# Comparison of the effects of metformin on MDA-MB-231 breast cancer cells in a monolayer culture and in tumour spheroids as a function of nutrient concentrations

**DOI:** 10.1101/351742

**Authors:** Maruša Bizjak, Petra Malavašič, Sergej Pirkmajer, Mojca Pavlin

**Author notes:** Corresponding author (MP). These authors contributed equally to this work (Maruša Bizjak, Petra Malavašič).

## Abstract

Metabolic pathways of cancer cells depend on the concentrations of nutrients in their micro-environment. However, they can vary also between monolayer cultures of cancer cells and tumour spheroids. Here we examined whether the absence of glucose, pyruvate and glutamine increases the sensitivity of MDA-MB-231 cells to metabolic drug metformin using two *in vitro* cell models (monolayer culture and tumour spheroids). To evaluate the effects of nutrient depletion in more detail, we tested the effects of metformin in commonly used media (DMEM, MEM and RPMI-1640) that differ mainly in the concentrations of amino acids. We used MTS, Hoechst and propidium iodide assay to determine cell number, viability and survival, respectively. We evaluated the effects of metformin on the size of tumour spheroids and determined cell survival by calcein and propidium iodide staining. Finally, we observed the effects of metformin in nutrient depleted conditions on the phosphorylation of AMP-activated protein kinase using Western blotting. Our main finding is that the effects of metformin on MDA-MB-231 cells depend on *in vitro* cell model used (monolayer culture vs. tumour spheroids). While metformin did not have any major effect on proliferation of MDA-MB-231 cells grown in complete cell culture media in a monolayer culture, it disintegrated tumour spheroids in MEM and RPMI-1640 medium. The effects of metformin on tumour spheroids were most pronounced in MEM, which is deficient of several non-essential amino acids. Glutamine depletion had no effect on the sensitivity of MDA-MB-231 cells to metformin in all tested conditions, whereas pyruvate depletion sensitized MDA-MB-231 cells to metformin in a monolayer culture only in MEM. Taken together, our results show that media formulation as well as *in vitro* cell model (monolayer culture vs. tumour spheroids) must be considered, when we evaluate the effects of metformin on MDA-MB-231 cells as a function of nutrient availability.

## Introduction

Triple negative breast cancer, which lacks estrogen and progesterone receptors and is negative for human epidermal growth factor receptor 2 (HER-2), is highly aggressive form of breast cancer with limited treatment options [1]. Cancer cells have altered metabolic pathways in comparison to normal cells. The first metabolic alteration that was described in cancer cells is their increased consumption of glucose under aerobic conditions, which is also known as the Warburg effect [2]. Moreover, cancer cells differ from normal cells in consumption of other nutrients [3] as well as in intrinsic characteristics that affect metabolic pathways [4,5]. Metabolic phenotype of cancer cells at least partially depends on their micro-environment, which is often depleted of some nutrients [6–8]. Furthermore, under *in vitro* conditions metabolic pathways differ between 2D monolayer cultures of cancer cells and the physiologically more relevant 3D tumour spheroids [9–12]. Importantly, the type of *in vitro* cell model (monolayer culture vs. tumour spheroids) and media formulation might also alter the sensitivity of breast cancer cells to pharmaceutical compounds that target cell metabolism and has a potential to treat breast cancer. One of such compounds is metformin.

Metformin is the most commonly used oral drug to treat type 2 diabetes and has potential anti-cancer effects in patients with breast cancer [13–15]. However, its mechanism of action is still not completely understood. In type 2 diabetes, metformin ameliorates glucose homeostasis and alleviates hyperinsulinemia, thus reducing the risk factors for development of insulin-sensitive cancers [16]. Besides its systemic action, metformin can also target cancer cells directly. Its main direct mechanism of action is probably suppression of cellular oxidative phosphorylation via inhibition of complex I in mitochondrial respiratory chain [17–20]. Another important mechanism is inhibition of mitochondrial glycerophosphate dehydrogenase, which reduces gluconeogenesis in hepatocytes [21]. Metformin-mediated inhibition of mitochondrial oxidative phosphorylation reduces cellular NAD+/NADH ratio [22,23], attenuates mitochondrial anaplerotic reactions [24,25], in particular aspartate biosynthesis [22,23], and induces reductive metabolism of glutamine-derived carbon in tricarboxylic acid (TCA) cycle [24,26,27]. Furthermore, metformin activates AMP-activated protein kinase (AMPK), which is the main regulator of cellular energy homeostasis [28]. Once activated, AMPK accelerates catabolic and inhibits anabolic processes in cancer cells, which enables survival of cancer cells in energetic crisis [29,30]. Metformin-stimulated AMPK activation in breast cancer cells is more pronounced in glucose depleted conditions [30,31], while AMPK does not mediate the effects of metformin under serine depletion [33]. The effects of depletion of other nutrients on metformin-stimulated AMPK activation are still largely unknown.

Cancer cell sensitivity to metformin *in vitro* is not merely an intrinsic property of cancer cells, but can be modulated by nutrient concentrations in cell culture media [17,22,23,31–36] and consequently also by medium renewal protocols [31]. Metformin suppresses proliferation of various cancer cells more effectively in medium without glucose [31,36–39] or pyruvate [22,23,33,34], while the absence of glutamine increases metformin’s effects on liver cancer cells [26]. Studies imply that the absence of several amino acids might increase the effects of metformin on cancer cells synergistically with the absence of other nutrients [22,33,35]. However, the effects of depletion of nutrient combinations on the sensitivity of the most wieldy used triple negative breast cancer cells, MDA-MB-231 cells [40], to metformin were not examined in detail. Besides, although it is well-known that metformin affects cancer cells grown in various 3D cell culture models [41–45], the role of nutrient availability on the effects of metformin on tumour spheroids remains unknown.

To our knowledge, the effects of metformin on MDA-MB-231 cells as a function of nutrient concentrations have never been directly compared between a 2D monolayer cell culture and 3D tumour spheroids. Here we examined whether the absence of three major nutrients, glucose, pyruvate and glutamine, increases the sensitivity of MDA-MB-231 cells to metformin in a monolayer culture and in tumour spheroids. We tested effects of nutrients and metformin using three widely used media (DMEM, MEM and RPMI-1640) that differ mainly in the concentrations of amino acids. Our main finding is that sensitivity of MDA-MB-231 cells to metformin varies between a monolayer culture and the more physiologically relevant tumour spheroids. Furthermore, usage of MEM increased the effects of metformin on tumour spheroids and sensitized MDA-MB-231 cells in a monolayer culture to metformin in pyruvate-depleted condition.

## Materials & Methods

### Antibodies and reagents

Antibodies against phospho-ACC (Ser^79^) (CST3661), and phospho-AMPKα (Thr^172^) (CST2535) were form Cell Signaling Technology. Antibodies against actin were from Cell Signaling Technology. Bis-Tris 4–20% polyacrylamide gels were from Sigma-Aldrich. TruPAGE™ TEA-Tricine SDS Running Buffer was from Sigma-Aldrich and horseradish peroxidase secondary antibody conjugate (170–6515) were from Bio-Rad. Polyvinylidene Fluoride (PVDF) Immobilon-P membrane (IPVH00010) was from Merck and protein molecular weight marker (P7712S) from New England BioLabs. Enhanced chemiluminescence (ECL) reagent was from Life Technologies (Thermo Fisher Scientific). Metformin was from Calbiochem (Merck Millipore). All other reagents, unless otherwise specified, were from Sigma-Aldrich.

### MDA-MB-231 cell culture

MDA-MB-231 cells were from ATCC (USA). MDA-MB-231 cells were grown in RPMI-1640 medium (Genaxxon bioscience, Germany) supplemented with 4.5 g/l of glucose, 2 mM L-glutamine, 1 mM pyruvate and 10 % fetal bovine serum (FBS; Sigma-Aldrich). They were maintained at 37 °C in a humidified atmosphere with 5% (v/v) CO_2_. Experiments were performed in RPMI-1640 (Genaxxon, custom made medium) or DMEM (Gibco, A14430) with or without 4.5 g/l of glucose, 2 mM glutamine and 1 mM pyruvate. Alternatively, some experiments were performed in MEM medium (Sigma, M5775). MEM medium was supplied with 1 g/l of glucose so we increased glucose concentration of MEM in all the experiments to match RPMI and DMEM medium (4.5 g/l). Furthermore, MEM medium was also supplemented with 2 mM glutamine and/or 1 mM pyruvate. Cell culture media used differ mainly in the concentrations of amino acids (Table 1).

**Table 1.** The concentrations of amino acids in cell culture media.

| Amino acids | RPMI-1640 | DMEM | MEM | RPMI-1640 | DMEM | MEM |
| --- | --- | --- | --- | --- | --- | --- |
|  | mM |  |  | Relative to RPMI-1640 |  |  |
| L-Arginine hydrochloride | 1.15 | 0.398 | 0.597 | 1.00 | 0.346 | 0.519 |
| L-Asparagine × H<sub>2</sub>O | 0.333 | 0 | 0 | 1.00 | 0 | 0 |
| L-Aspartic acid | 0.150 | 0 | 0 | 1.00 | 0 | 0 |
| L-Cystine | 0.208 | 0.201 | 0.0999 | 1.00 | 0.967 | 0.480 |
| L-Glutamic acid | 0.136 | 0 | 0 | 1.00 | 0 | 0 |
| Glycine | 0.133 | 0.400 | 0 | 1.00 | 3.00 | 0 |
| L-Histidine hydrochloride-H<sub>2</sub>O | 0.0967 | 0.200 | 0.200 | 1.00 | 2.07 | 2.07 |
| L-Hydroxyproline | 0.153 | 0 | 0 | 1.00 | 0 | 0 |
| L-Isoleucine | 0.382 | 0.801 | 0.397 | 1.00 | 2.10 | 1.04 |
| L-Leucine | 0.382 | 0.801 | 0.397 | 1.00 | 2.10 | 1.04 |
| L-Lysine hydrochloride | 0.219 | 0.798 | 0.396 | 1.00 | 3.65 | 1.81 |
| L-Methionine | 0.101 | 0.201 | 0.101 | 1.00 | 2.00 | 1.00 |
| L-Phenylalanine | 0.0909 | 0.400 | 0.194 | 1.00 | 4.40 | 2.13 |
| Proline | 0.174 | 0 | 0 | 1.00 | 0 | 0 |
| L-Serine | 0.286 | 0.400 | 0 | 1.00 | 1.40 | 0 |
| L-Threonine | 0.168 | 0.798 | 0.403 | 1.00 | 4.75 | 2.40 |
| L-Tryptophan | 0.0245 | 0.0784 | 0.049 | 1.00 | 3.20 | 2.00 |
| Tyrosine | 0.110 | 0.398 | 0.230 | 1.00 | 3.61 | 2.09 |
| L-Valine | 0.171 | 0.803 | 0.393 | 1.00 | 4.70 | 2.30 |
| Total amino acids | 4.47 | 6.68 | 3.46 | 1.00 | 1.49 | 0.774 |

### MTS cell viability assay

MTS assay (Promega Corp, Fitchburg, WI, USA) was performed as described previously[31]. Briefly, upon completion of the experiment, MDA-MB-231 cells were washed with phosphate saline buffer (PBS) and placed in a serum-free RPMI-1640 medium with 4.5 g/l of glucose, 2 mM glutamine and 1 mM pyruvate. Then, Cell Titer®96AQueous One (MTS) solution (Promega Corp, Fitchburg, WI, USA) was added to each well and cells were incubated for about 1-hour at 37 °C, 5 % CO_2_. Absorbance of supernatant was measured at 490 nm using the Tecan Infinite 200 (Tecan Group Ltd, Männedorf, Switzerland).

### Cell number

The number of MDA-MB-231 cells was determined as described previously[31,36]. Briefly, medium was removed from each well and plates were frozen at −20 °C. On the day of the analysis, cells were thawed and lysed with a 0.04% SDS solution at room temperature for 30 minutes. Then, buffer containing 50 mM TRIS-HCl, 100 mM NaCl (pH = 8.25) and 5 μg/ml Hoechst 33342 stain (Thermo Fisher Scientific) was added to each well. Fluorescence intensity was determined at 350 nm excitation and 461 nm emission using Tecan Infinite 200 (Tecan, Männedorf, Switzerland).

### Propidium iodide staining

MDA-MB-231 cells were seeded in 12-well plates. Next day, they were placed in a fresh cell culture medium and treated with 5 mM metformin. At the end of treatment, MDA-MB-231 cells were collected, pelleted and resuspended in PBS. Propidium iodide was added to a final concentration of 0.15 mM and cell suspension was analysed by Attune™ NxT flow cytometer (Thermo Fisher Scientific, Waltham, USA). Propidium iodide signal of at least 2 × 10^4^ events per sample was collected using the BL-2 filter (574/26). Final analysis was performed with Attune^®^ Cytometric Software (Thermo Fisher Scientific, Waltham, USA).

### Tumour spheroids

For formation of 3D cellular spheroids, MDA-MB-231 cells were seeded in various cell culture media in U-shaped low adherent 96 well cell culture plates (Corning, New York, USA). Following seeding, plates were centrifuge for 2 minutes at 300 rcf. After 72 hours, tumour spheroids were stained with calcein (1 μM final concentration, Life Technology) and propidium iodide (0.15 mM final concentration) for 10 and 5 minutes, respectively. Images were acquired using fluorescent Leica DM IL LED microscope (Leica Microsystem) and analyzed by ImageJ Software (National Institute of Helath, USA).

### Western blotting

MDA-MB-231 cells were treated with 5 mM metformin in complete RPMI-1640 medium or in RPMI-1640 medium without glutamine, pyruvate or glucose for 24 hours. Then cells were washed twice with ice-cold PBS and harvested in Laemmli buffer (62.5 mM Tris-HCl, pH 6.8, 2% (w/v) sodium dodecyl sulfate (SDS), 10% (v/v) glycerol, 5% 2-mercaptoethanol, 0.002% bromophenol blue). Total protein concentration was measured by Pierce 660 (Thermofisher). Samples, containing equivalent amount of proteins, were loaded on a 4–20% polyacrylamide gel (TruPAGE™ Precast Gels, Sigma) and separated using electrophoresis (Mini-protean tetra cell system, Bio Rad). Subsequently, proteins were transferred to PVDF membrane. Ponceau S (0.1% (w/v) Ponceau S in 5% (v/v) acetic acid) was used to evaluate the efficiency of the protein transfer and sample loading. Membranes were blocked in 5% (w/v) skimmed milk in TBS-T (20 mM Tris, 150 mM NaCl, 0.02% (v/v) Tween-20, pH 7.5), which was followed by overnight incubation in primary antibodies at 4°C. After washing, membranes were incubated with the appropriate secondary horseradish peroxidase-conjugated antibody. Enhanced chemiluminescence using Agfa X-ray film was used to detect immuno-reactive. ImageJ was used for densitometric analysis.

### Statistical analysis

Statistical analysis was performed with GraphPad Prism (v6; GraphPad Software, Inc., La Jolla, CA, USA) using one-way ANOVA or two-way ANOVA, followed by Bonferroni, Tukey or Sidak test. Statistically significant results are displayed as follows: *P≤0.05; **P≤0.01, *** P≤0.001.

## Results

### The effect of nutrient availability on proliferation of MDA-MB-231 cells in RPMI-1640 medium

Nutrient availability in the microenvironment of cancer cells modulates their metabolism [3], while intrinsic properties of cancer cells determine their dependency on specific nutrients for proliferation and survival [4,5]. To assess nutrient requirements of MDA-MB-231 cells, we grew them in the complete RPMI-1640 medium or in the nutrient-deficient RPMI-1640 medium without pyruvate, glucose or glutamine for 72 hours (Fig. 1). In all our experiments complete medium contained 2 mM glutamine, 1 mM pyruvate and 4.5 g/l of glucose. The complete and nutrient-deficient media, lacking glucose, glutamine and/or pyruvate, were supplemented with 10% FBS. The depletion of glutamine and pyruvate did not affect MDA-MB-231 cell proliferation, whereas depletion of glucose slightly suppressed proliferation of cells after 72 hours (Fig. 1A). As a positive control for suppression of proliferation, we used serum-free RPMI-1640 medium, which blocked MDA-MB-231 cell proliferation (Fig. 1B). Thus, unlike depletion of serum growth factors, removal of glutamine and pyruvate does not have any major effect on proliferation of MDA-MB-231 cells.

**Figure 1:**
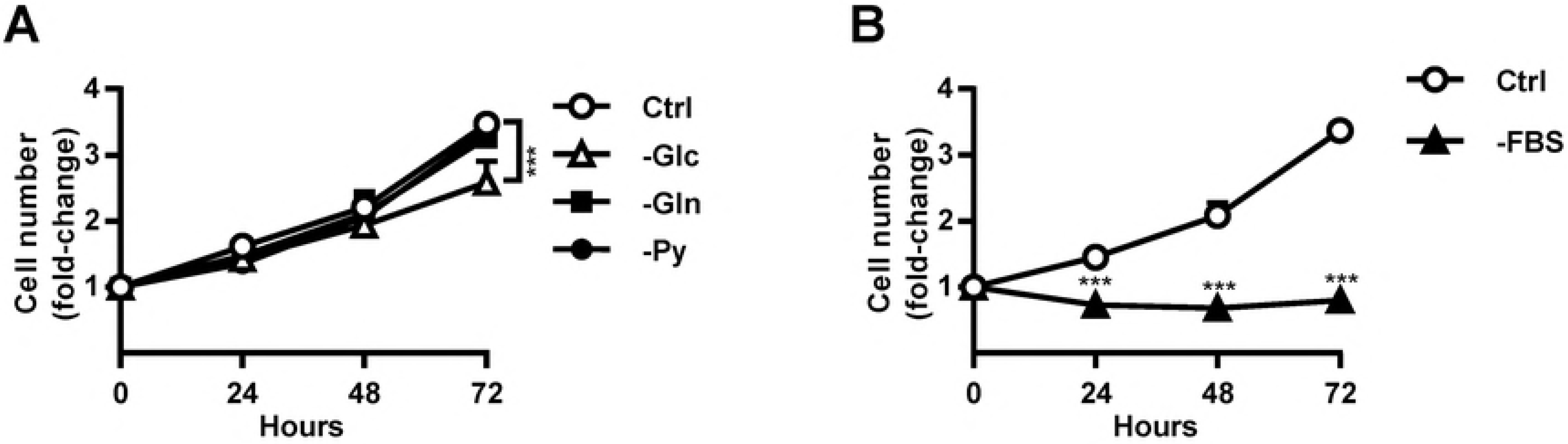
The effect of nutrient availability on proliferation of MDA-MB-231 cells in RPMI-1640 medium. MDA-MB-231 cells were grown for 72 hours in complete RPMI-1640 medium or in RPMI-1640 medium without glucose, glutamine or pyruvate (A). Alternatively MDA-MB-231 cells were grown for 72 hours in complete RPMI-1640 medium or in serum-free RPMI-1640 medium (B). Relative number of cells was determined by Hoechst staining. Results are means±SEM (n = 3-4).

### Nutrient availability determines the sensitivity of MDA-MB-231 cells to metformin

Limited availability of glucose and/or other nutrients increases the sensitivity of cancer cells to metformin [17,22,23,26,31–39,46,47]. However, excepting glucose, the role of other nutrients and their combinations on the sensitivity of MDA-MB-231 cells to metformin was not examined in detail. To determine whether depletion of pyruvate and glutamine affects sensitivity to metformin, we grew MDA-MB-231 cells in RPMI-1640 medium in the absence of glucose, pyruvate or glutamine and treated them with 5 mM metformin for 72 hours (Fig. 2). We determined the number and viability of MDA-MB-2321 cells using Hoechst staining and MTS assay [31], respectively (Fig. 2A-D). Metformin did not affect the number and viability of MDA-MB-231 cells grown in the complete RPMI-1640 medium or in RPMI-1640 medium lacking glutamine or pyruvate, while it significantly reduced the number and viability of MDA-MB-231 cells in RPMI-1640 medium without glucose (Fig. 2A, B).

**Figure 2:**
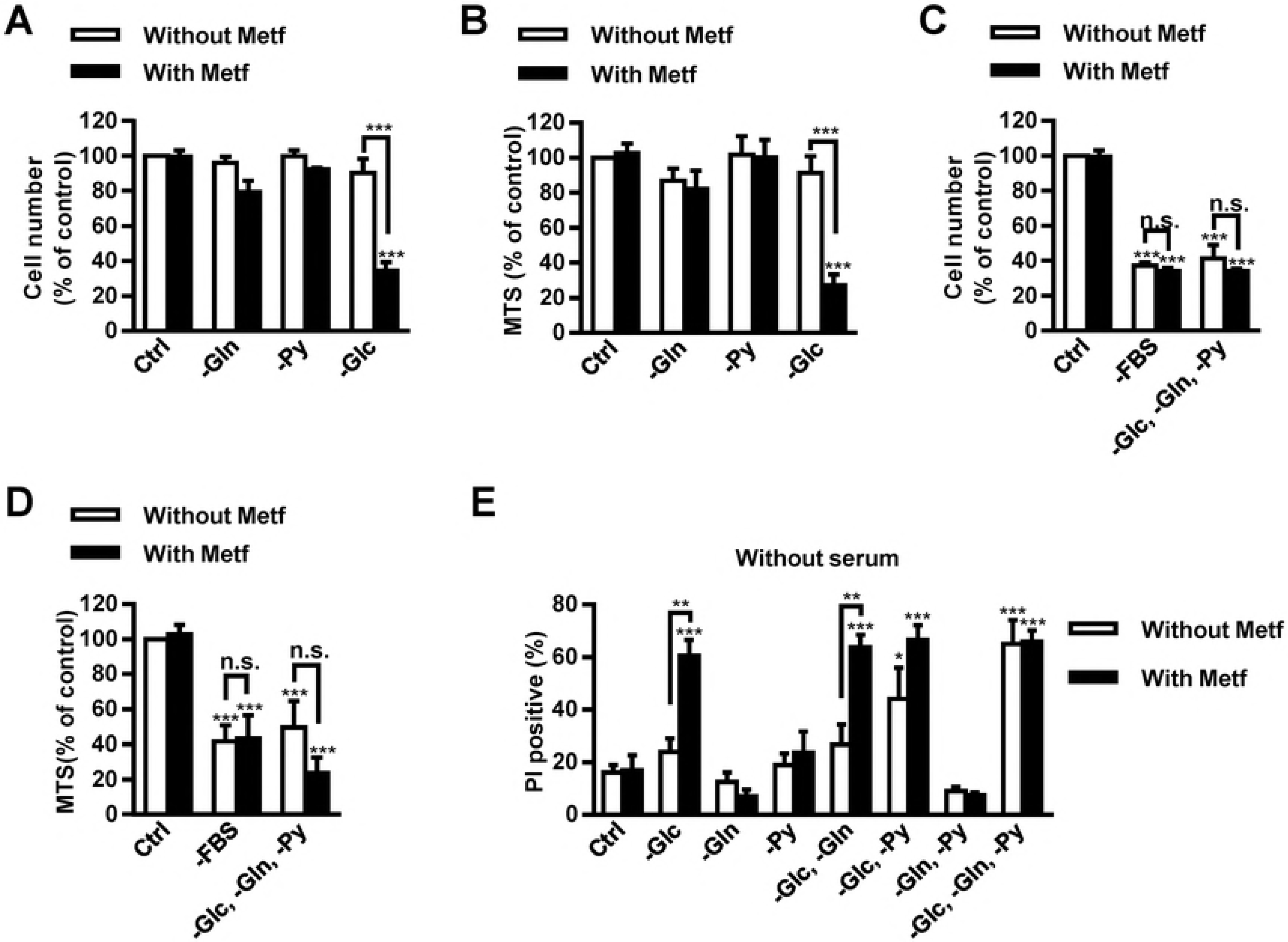
Nutrient availability determines the sensitivity of MDA-MB-231 cells to metformin. (A, B) MDA-MB-231 cells were grown in complete RPMI-1640 medium or in RPMI-1640 medium without glucose, glutamine or pyruvate and treated with 5 mM metformin. After 72 hours, cell number (A) and viability (B) were determined by Hoechst and MTS staining, respectively. Results are means±SEM (n = 2-4). (C, D) MDA-MB-231 cells were grown in complete RPMI-1640 medium, serum-free RPMI-1640 or in RPMI-1640 medium lacking glucose, glutamine and pyruvate (with serum) and treated with 5 mM metformin. After 72 hours, cell number (C) and viability (D) were determined by Hoechst and MTS staining, respectively. Results are means±SEM (n = 3-4). (E) MDA-MB-231 cells were cultured for 72 hours in serum-free RPMI-1640 medium without glutamine, glucose and/or pyruvate and treated with 5 mM metformin. Cell survival was determined by propidium iodide assay using flow cytometry. Results are means±SEM (n = 3-4).

Serum starvation causes changes in AMPK activation [48], indicating it might also affect the sensitivity of MDA-MB-231 cells to metformin, which is an indirect AMPK activator. To evaluate whether serum starvation or deficiency of all three analyzed nutrients (glutamine, glucose and pyruvate) affect sensitivity to metformin, we grew MDA-MB-231 cells for 72 hours in the complete RPMI-1640 medium, in serum-free RPMI-1640 medium (with glucose, glutamine and pyruvate) or in a nutrient-deficient RPMI-1640 medium with serum, but without glutamine, glucose and pyruvate (Fig. 2C, D). Serum-free RPMI-1640 medium and nutrient-deficient RPMI-1640 medium with serum markedly reduced the number of cells and their viability. Metformin did not further decrease the number or viability of MDA-MB-231 cells.

To determine whether nutrients and/or metformin affects survival of MDA-MB-231 in the absence of serum and specific nutrients, we cultured them in serum-free RPMI-1640 medium without glutamine, glucose and/or pyruvate (Fig. 2E). After 72-hour treatment with 5 mM metformin, we determined the survival of MDA-MB-231 cells using propidium iodide assay and flow cytometry. Compared with control, cell survival was reduced in RPMI-1640 medium lacking glucose and pyruvate and in RPMI-1640 medium lacking all three nutrients, glucose, pyruvate and glutamine. Metformin did not further reduce survival of cells grown under these two conditions. In comparison to non-treated cells, metformin reduced survival of cells grown in RPMI-1640 medium without glucose or in RPMI-1640 medium lacking glucose and glutamine.

### The size of tumour spheroids is increased by metformin and depends on nutrient availability

Metabolic phenotype of cancer cells depends on whether they are grown in 2D monolayer cultures or in 3D tumour spheroids [9–12]. We therefore evaluated if sensitivity to metformin differs between MDA-MB-231 cells grown in a monolayer culture and those grown in tumour spheroids. To this end, tumour spheroids were treated with 5 mM metformin in complete RPMI-1640 medium without glucose, glutamine or pyruvate for 72 hours (Fig. 3). We determined the size of each tumour spheroid and survival of MDA-MB-231 cells composing it, using calcein and propidium iodide staining. Metformin slightly increased the size of tumour spheroids in all tested conditions and spheroids became less compact. The effect was most pronounced in RPMI-1640 medium without glucose (Fig. 3A, B). Survival of metformin-treated MDA-MB-231 cells was decreased only in the absence of glucose (Fig 3C).

**Figure 3:**
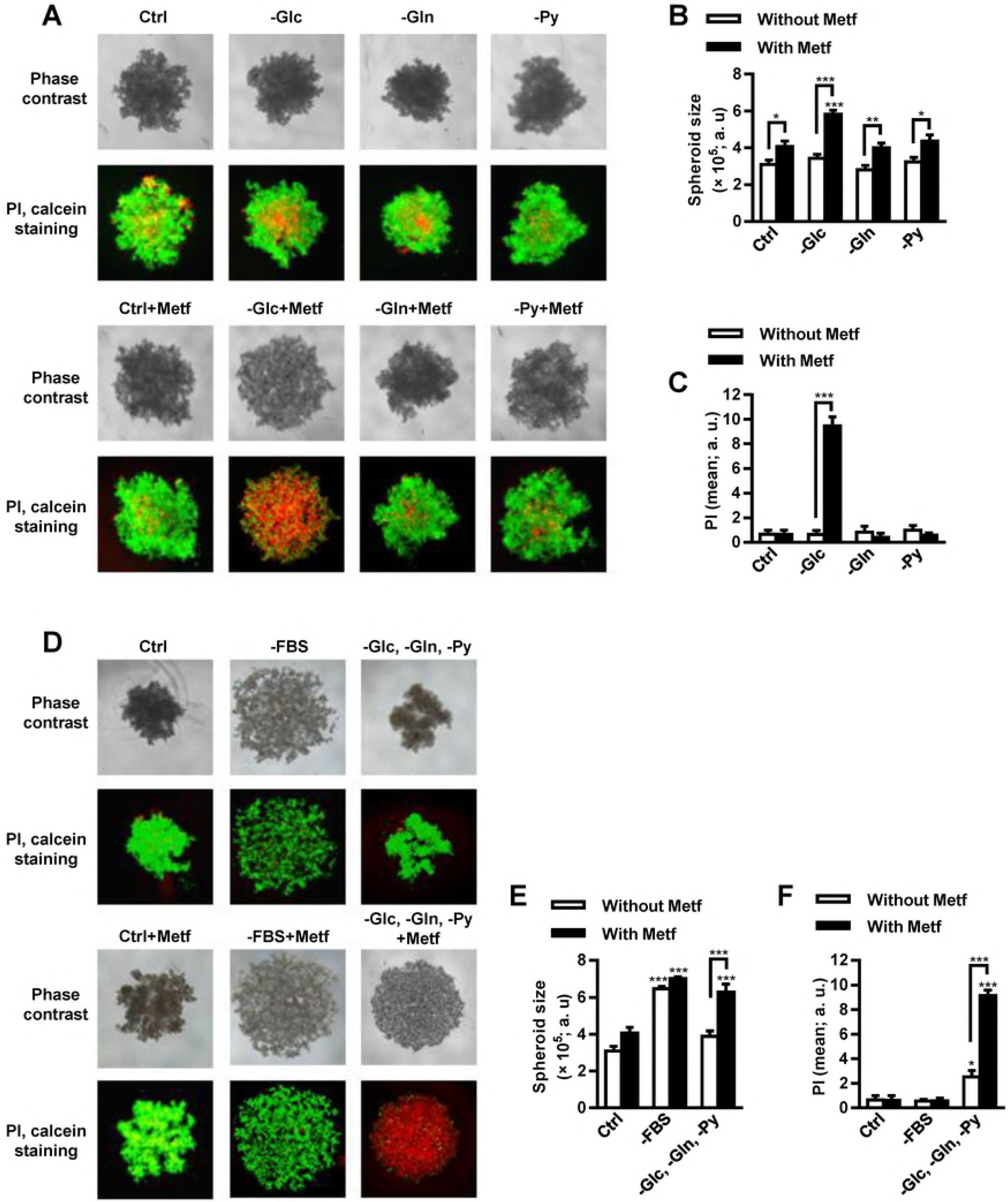
The size of tumour spheroids is increased by metformin and depends on nutrient availability. (A, B, C) Tumour spheroids were grown in complete RPMI-1640 medium or in RPMI-1640 medium without glucose, glutamine or pyruvate and treated with 5 mM metformin for 72 hours. Tumour spheroids were double stained by propidium iodide and calcein and observed using fluorescence microscopy (A) The size of tumour spheroids was determined by ImageJ programme. Results are means±SEM (n = 5). (B). Fluorescence intensity of propidium iodide positive MDA-MB-231 cells in each tumour spheroid was determined by ImageJ programme. Results are means±SEM (n = 3) (C). (D, E, F) Tumour spheroids were grown in complete RPMI-1640 medium, in serum-free RPMI-1640 medium or in RPMI-1640 medium lacking glucose, glutamine and pyruvate (with serum). Tumour spheroids in all conditions were treated with 5 mM metformin for 72 hours. Tumour spheroids were double stained by propidium iodide and calcein and observed using fluorescence microscopy (D). Size 14 of tumour spheroids was determined by ImageJ programme. Results are means±SEM (n = 3-5) (E). Fluorescence intensity of propidium iodide positive MDA-MB-231 cells in each tumour spheroid was determined by ImageJ programme. Results are means±SEM (n = 2-3) (F).

We also investigated the effects of serum-free RPMI-1640 medium and the absence of all three nutrients, glucose, glutamine and pyruvate, on the size and survival of tumour spheroids treated with 5 mM metformin for 72 hours. Tumour spheroids grown in serum-free RPMI-1640 medium with or without 5 mM metformin were completely disintegrated (Fig. 3D, E). Metformin did not significantly affect the percentage of dead cells composing tumour spheroids in this condition. On the other hand, 5 mM metformin increased the size of tumour spheroids and the percentage of dead cells composing them in RPMI-1640 medium without glucose, glutamine and pyruvate (Fig. 3D, F).

### Metformin disintegrates tumour spheroids grown in MEM

Formulation of cell culture medium affects the sensitivity of cancer cells to metformin in a monolayer culture [22,35], while the effects of nutrient availability on the sensitivity of MDA-MB-231 cells in tumour spheroids remain largely unknown. We therefore compared the effects of metformin on MDA-MB-231 cells grown in tumour spheroids in RPMI-1640 medium with its effects in DMEM and MEM that are also commonly used in cancer cell culturing. DMEM has higher concentrations of the majority of amino acids than RPMI-1640 medium, but does not contain aspartate. MEM does not contain non-essential amino acids: alanine, glycine, glutamate, proline, serine, asparagine and aspartate (Table 1).

Tumour spheroids were grown in complete DMEM or MEM and treated with 5 mM metformin for 72 hours. Cell survival was determined by propidium iodide and calcein staining (Fig. 4). Untreated spheroids had similar size independent from cell culture medium. Metformin disintegrated tumour spheroids grown in MEM but not in DMEM (Fig. 4A, B). Cell survival was lower in untreated tumour spheroids grown in MEM than in DMEM. Metformin did not further reduce cell survival in tumour spheroids grown in MEM or DMEM (Fig. 4C).

**Figure 4:**
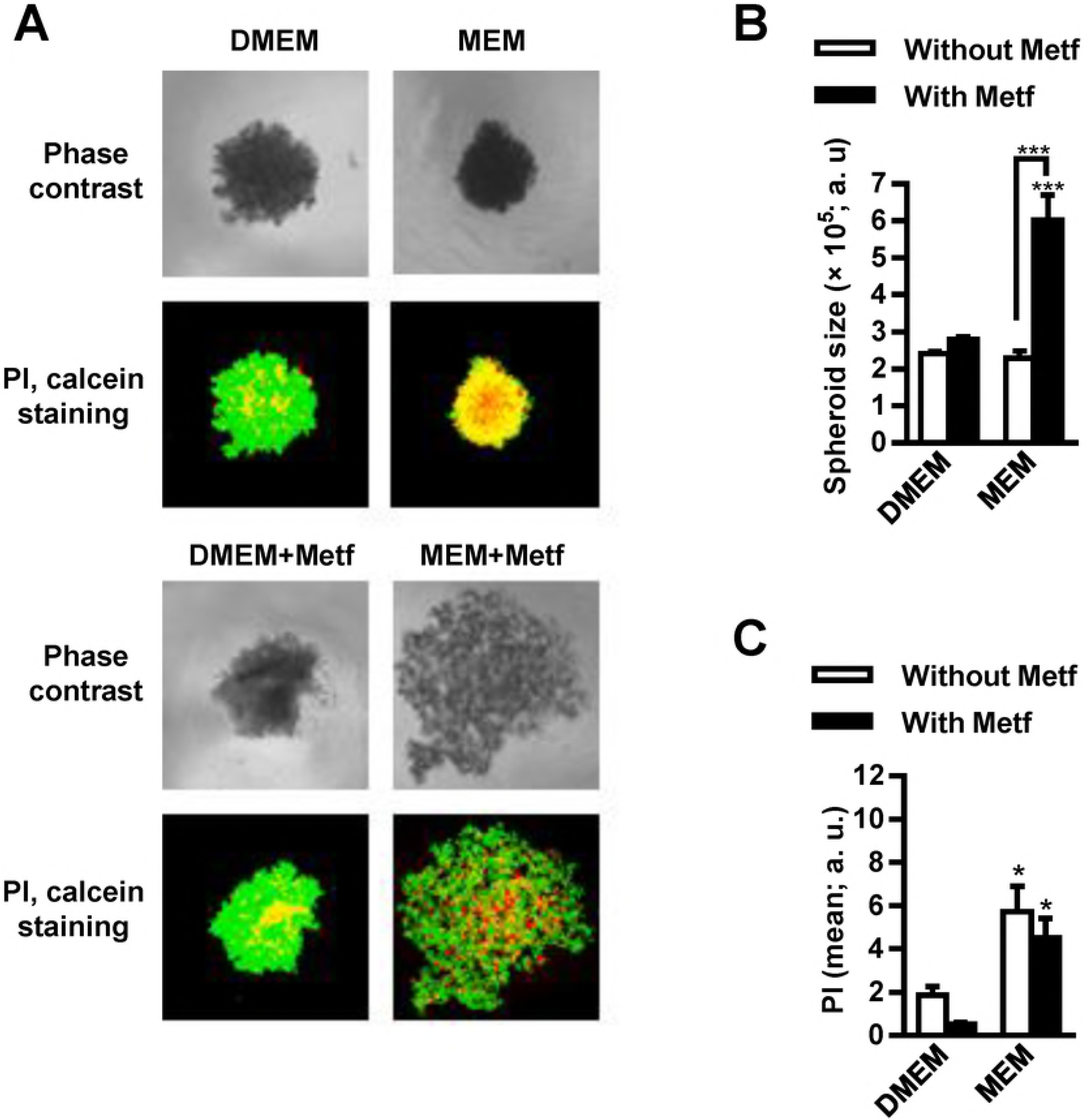
Metformin disintegrates tumour spheroids grown in MEM. Tumour spheroids were grown in complete DMEM or MEM and treated with 5 mM metformin for 72 hours. Tumour spheroids were double stained by calcein and propidium iodide and observed by fluorescence microscopy (A). Relative size of tumour spheroids was determined by ImageJ programme. Results are means±SEM (n = 3) (B). Fluorescence intensity of propidium iodide positive cells was determined by ImageJ programme Results are means±SEM (n = 3) (C).

### Effects of nutrient availability on AMPK activation

Glucose depletion increases AMPK activation by metformin [31,32]. Whether depletion of other nutrients has similar effects on the sensitivity of MDA-MB-231 cells to metformin has not been examined in detail. Thus, we compared the effects of deficiency of pyruvate, glutamine and/or glucose on metformin-stimulated AMPK activation in MDA-MB-231 cells (Fig. 5). Activation of AMPK was estimated by measuring phosphorylation of AMPK (Thr^172^) and its downstream target acetyl-CoA carboxylase (ACC). We treated MDA-MB-231 cells with 5 mM metformin in DMEM without glutamine, pyruvate and/or glucose for 24 hours. Metformin increased phosphorylation of AMPK in DMEM without glucose and in DMEM without glucose and pyruvate (Fig. 5A, B). Metformin increased phosphorylation of ACC in the absence of pyruvate, glucose and their combination (Fig. 5A, C). Deficiency of glutamine did not enhance metformin-stimulated AMPK or ACC phosphorylation.

**Figure 5:**
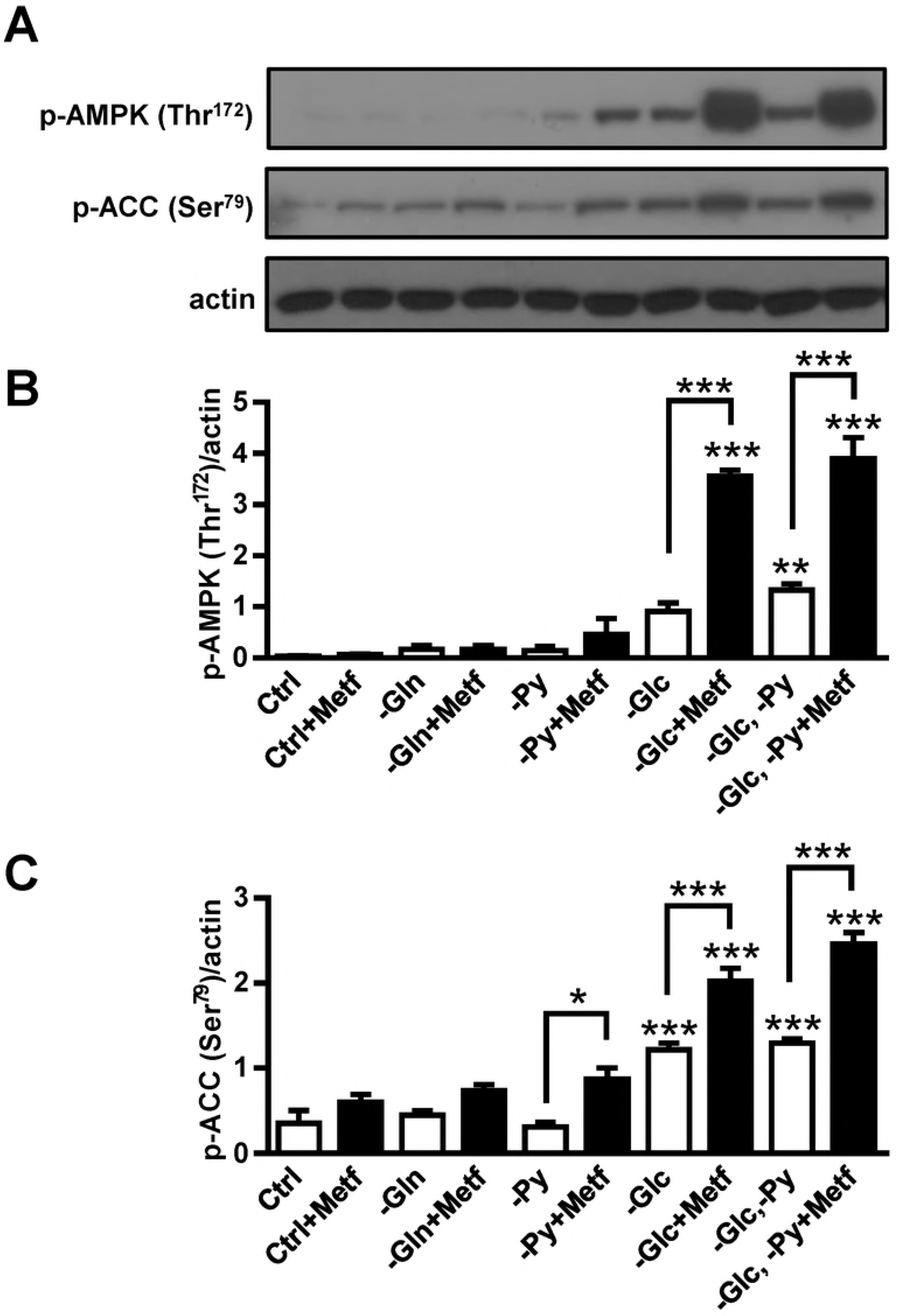
Effects of nutrient availability on AMPK activation. MDA-MB-231 cells were grown in DMEM without glutamine, glucose and/or pyruvate and treated with 5 mM metformin for 24 hours. Phosphorylation of AMPK (Thr^172^) (A, B) and phosphorylation of ACC (Ser^79^) (C) was measured by Western blot. Results are means±SEM (n = 3).

### Metformin reduces the number of MDA-MB-231 cells grown in MEM without pyruvate

Cell culture media contain various concentrations of amino acids (Table 1), which might alter the sensitivity of MDA-MB-231 cells to metformin in the absence of glutamine, pyruvate and/or glucose. We tested how depletion of glucose, glutamine and pyruvate affects proliferation and survival of MDA-MB-231 cells grown in DMEM and MEM (Fig. 6). MDA-MB-231 cells were grown in a complete DMEM lacking glutamine, pyruvate and/or glucose and treated with 5 mM metformin for 96 hours. Metformin did not affect proliferation and survival of MDA-MB-231 cells grown in the complete DMEM and DMEM without glutamine or pyruvate (Fig. 6A, B). However, it suppressed proliferation of MDA-MB-231 cells and reduced their survival in DMEM medium without glucose. The effect of metformin on cell survival was even more profound when DMEM was without both glucose and pyruvate (Fig. 6B).

**Figure 6:**
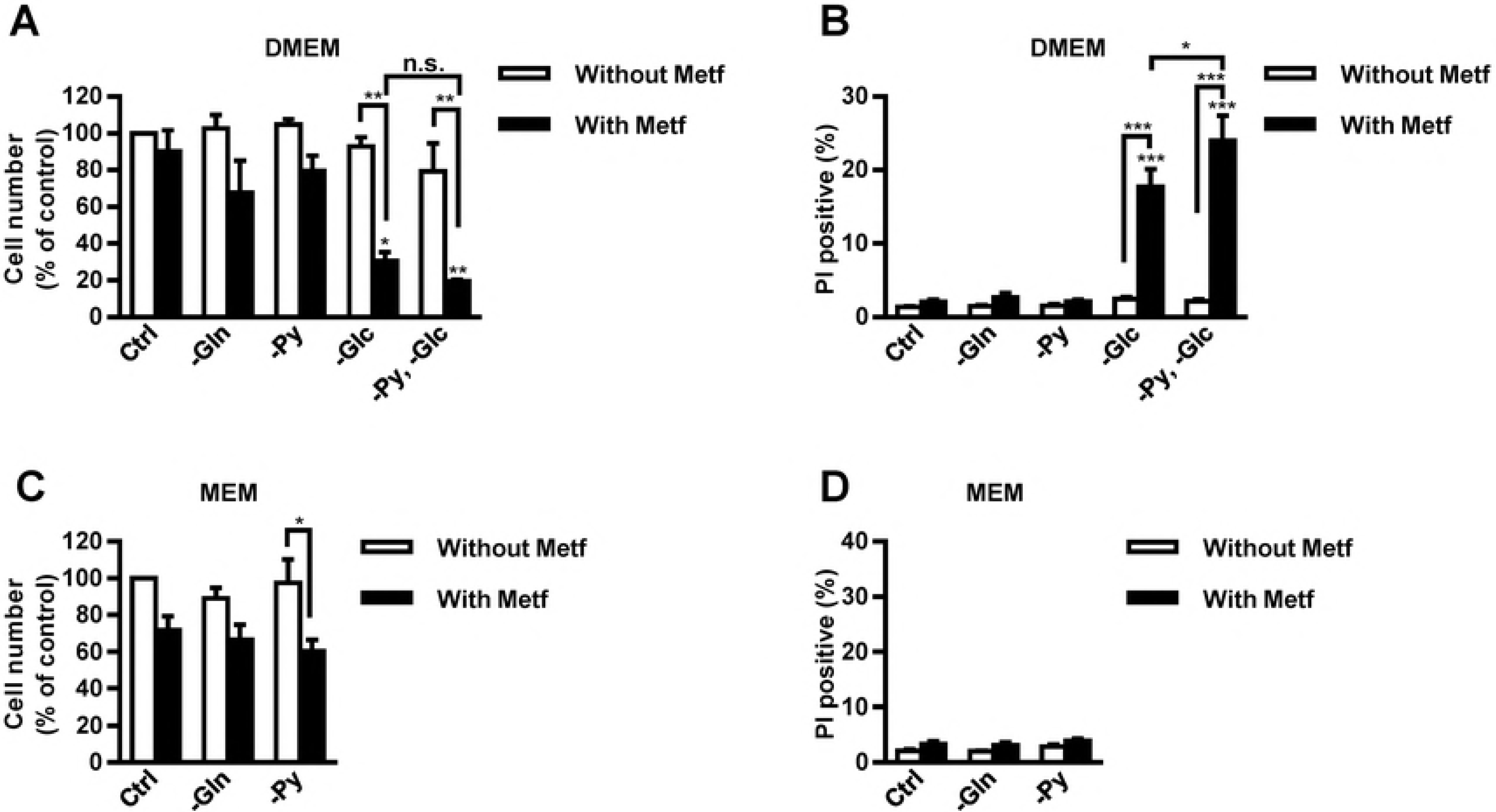
Metformin reduces the number of MDA-MB-231 cells grown in MEM without pyruvate. (A, B) MDA-MB-231 cells were grown in complete DMEM or in DMEM without glutamine, pyruvate and/or glucose and treated with 5 mM metformin for 96 hours. Relative cell number (A) and propidium iodide positive cells (B) were determined by Hoechst and propidium iodide staining, respectively. Results are means±SEM (n = 3). (C, D) MDA-MB-231 cells were grown in complete MEM or in MEM without glutamine or pyruvate and treated with 5 mM metformin for 96 hours. Relative cell number (C) and propidium iodide positive cells (D) were determined by Hoechst and propidium iodide staining, respectively. Results are means±SEM (n = 2-3).

In the absence of metformin, depletion of glutamine or pyruvate did not affect survival and the number of MDA-MB-231 cells in MEM (Fig. 6C, D). Metformin treatment did not significantly reduce the number and survival of cells grown in the complete MEM or in MEM without glutamine. In contrast, metformin reduced the number of MDA-MB-231 cells grown in MEM without pyruvate to about 60% (Fig. 6C), but it did not have any direct effect on their survival (<4% propidium iodide positive MDA-MB-231 cells) (Fig. 6D).

## Discussion

The metabolic phenotype of cancer cells is partially determined by nutrient concentrations in their microenvironment [3]. Metabolic pathways of cancer cells *in vitro* vary also between monolayer cultures and physiologically more relevant tumour spheroids [9–12]. Nutrient availability and metabolic phenotype can modify the sensitivity of cancer cells to pharmacological compounds that target cancer cell metabolism, such as metformin [17,22,23,26,31–39,46,47]. Recently, we have shown that maintaining constant glucose concentrations by daily medium renewal blocks the effects of metformin on MDA-MB-231 cells [31]. In contrast, MDA-MB-231 cells are sensitive to metformin in glucose-depleted conditions [31,32,36–39,47]. In the present study, we compared the effects of three major nutrients, glucose, glutamine and pyruvate, on the sensitivity of MDA-MB-231 cells to metformin in a monolayer culture and in tumour spheroids. Previous studies have shown that the sensitivity of cancer cells to metformin in a monolayer culture depends on media formulation [22,35]. Importantly, the formulations of commonly used media vary greatly in the concentrations of several nutrients. Here we show that the effects of metformin on MDA-MB-231 cells as a function of nutrient concentrations depend on *in vitro* cell model used (monolayer culture vs. tumour spheroids). Furthermore, we show that media formulation modulates the sensitivity of MDA-MB-231 cells to metformin upon pyruvate depletion.

Although 5 mM metformin did not have any major effect on proliferation of MDA-MB-231 cells grown in complete cell culture media in a monolayer culture, it disintegrated tumour spheroids in MEM and RPMI-1640 medium. This is broadly consistent with other studies that observed the effects of metformin on various 3D cell culture models [41–45]. The effect of metformin on disintegration of tumour spheroids was greatest in MEM, which lacks alanine, glycine, glutamate, proline, serine, asparagine and aspartate (Table 1). Thus, the effect of metformin on cell-cell interactions in tumour spheroids is enhanced if medium is deficient in non-essential amino acids. Furthermore, poor penetration of amino acids, which are present in lower concentrations in MEM, might contribute to reduced survival of MDA-MB-231 cells in tumour spheroids. On the other hand, MDA-MB-231 cells in tumour spheroids grown in MEM might have reduced survival upon treatment with metformin, because they were unable to adapt to metabolic stress. For instance, they were unable to cope with increased glycolytic rate that was partially required as a response to inhibition of oxidative phosphorylation [39] and partially to restore energy homeostasis upon detachment and to resist anoikis, a cell death mechanism induced by a loss of cell-cell or cell-matrix interactions [49].Taken together, in order to better predict effects of metformin in tissues, it is useful to evaluate its effects in nutrient limited conditions in tumour spheroids, which better mimic micro-environment of cancer cells in tumors.

Combined treatment with metformin and inhibitors of glutaminase synergistically reduce survival of prostate cancer cells [26]. In addition, glutamine depletion increases sensitivity of Huh7 liver cancer cells to metformin [26], which indicates that the absence of glutamine might also sensitize MDA-MB-231 cells to metformin. However, we show that media lacking glutamine did not enhance effects of metformin on proliferation and survival of MDA-MB-231 cells and did not have any major effects on tumour spheroids in comparison to the effects of metformin in complete medium. Glutamine is an important source of nitrogen that is required for protein and nucleotide biosynthesis. Besides, several studies show that cells treated with metformin relay on reductive metabolism of glutamine to support TCA cycle [24,26,27]. Since metformin failed to reduce viability of MDA-MB-231 cells in media lacking glutamine, we may speculate that these cells produce sufficient amount of α-ketoglutarate and obtain enough nitrogen from other nutrient sources [50]. Metformin attenuates anaplerotic flux of glutamine-derived carbon into the TCA cycle [24], which together with our results indicates that glucose is probably the major carbon source to sustain proliferation of metformin-treated MDA-MB-231 cells [46]. Furthermore, upon metformin-mediated inhibition of oxidative phosphorylation [18,19], MDA-MB-231 cells might produce the vast majority of energy from glycolysis. Taken together, our results suggest that media replete with glutamine do not block action of metformin on MDA-MB-231 cells in tumour spheroids or in a monolayer cell culture.

Metformin suppressed proliferation of MDA-MB-231 cells and reduced their survival under glucose-depleted conditions. Metformin attenuates mitochondrial anaplerotic reactions [24] and concomitantly augments glucose consumption of cancer cells [24,46,51]. MDA-MB-231 cells treated with metformin in the absence of glucose are thus unable to cope with energetic stress, which leads to suppressed cell proliferation and ultimately to cell death [17,31,39]. Energetic stress[52] or glucose depletion [53] activates AMPK, which upregulates catabolic processes in the cell and is promoting cell survival [29,30]. Cancer cells with impaired AMPK activation pathway are more sensitive to phenformin, an analogue of metformin [54]. In addition, metformin activates metabolic pathways that suppress proliferation of cancer cells also in an AMPK-independent manner [24,55]. The impairment of AMPK activation pathway increases the sensitivity of lung and colon cancer cells to metformin in glucose-depleted condition [56], which suggests that metformin-stimulated AMPK activation in medium without glucose that was observed in our study has a prosurvival role. We used 5 mM metformin, which is more than 100-times higher concentration than the one observed in plasma of diabetic patients [57]. However, *in vitro* studies using millimolar concentrations of metformin can predict the effects of metformin on tumour tissue in a murine model that has similar plasma concentrations of metformin than diabetic patients [20,34,58,59]. Therefore, since metformin in the absence of glucose reduced survival of MDA-MB-231 cells grown in both *in vitro* cell models (monolayer culture and tumour spheroids) it would probably also reduce survival of poorly perfused cancer cells *in vivo,* such as those in the center of the tumour.

Pyruvate depletion attenuates proliferation of cells with impaired oxidative phosphorylation [60–62], which was recently demonstrated also for cancer cells treated with metformin [22,23,34]. However, we observed that metformin suppressed proliferation of MDA-MB-231 cells only in MEM without pyruvate, but not in pyruvate-depleted DMEM or RPMI-1640 medium. On the other hand, depletion of pyruvate and glucose synergistically reduced survival of metformin-treated MDA-MB-231 cells grown in DMEM. Combined depletion of both nutrients increased metformin-stimulated AMPK activation, while pyruvate depletion alone enhanced metformin-stimulated phosphorylation of ACC, a downstream target of AMPK. Recent studies indicate that metformin suppresses proliferation of cancer cells in an AMPK-independent manner [22,24], which suggests that AMPK activation in the absence of glucose and pyruvate is probably a pro-survival response of MDA-MB-231 cells to energetic stress [29,30]. Pyruvate as well as glucose can be metabolized to acetyl-CoA and oxidized in the TCA cycle, thus driving ATP production. However, since metformin attenuates mitochondrial anaplerotic reactions [24,25] and suppresses oxidative phosphorylation [18,19], pyruvate probably blocks the effects of metformin through mechanisms independent from ATP production. Metformin decreases the NAD+/NADH ratio in cancer cells [22,33]. Thus, one of the main roles of exogenous pyruvate in cells with impaired oxidative phosphorylation is to support NAD+ production via its conversion into lactate [22,62]. NAD+ drives glycolysis, which may explain the synergistic effects of pyruvate and glucose depletion on survival of metformin-treated MDA-MB-231 cells.

Second role of pyruvate in cells with impaired oxidative phosphorylation is to support aspartate synthesis [22,60,62]. The addition of aspartate prevents metformin-mediated suppression of proliferation of some cancer cells [22,23,34], indicating that this effect depends on their intrinsic properties and/or the availability of other nutrients. In our study metformin tended, but did not significantly suppress proliferation of MDA-MB-231 cells grown in DMEM without pyruvate, which did not contain aspartate (Table 1). Breast cancer cells can produce pyruvate from glucose as well as from serine and glycine [63]. Besides, the absence of serine suppresses glycolysis [64] and thus prevents an adaptive response to inhibition of oxidative phosphorylation [33]. DMEM without pyruvate contains serine and glycine, which can block the effects of metformin. Indeed, metformin suppressed proliferation of MDA-MB-231 cells in MEM without pyruvate, which does not contain several nonessential amino acids, including aspartate, serine and glycine (Table 1). Taken together, our results show that metformin-treated MDA-MB-231 cells require pyruvate, which can be acquired per se or derived from glucose, serine and glycine, for their optimal proliferation. Thus, medium formulation must be considered, when we evaluate the effects of pyruvate-depletion on MDA-MB-231 cells.

## Conclusions

Here we show that the effects of metformin on MDA-MB-231 cells depend on *in vitro* cell model used (monolayer culture vs. tumour spheroids). While metformin had no effects on MDA-MB-231 cells in a monolayer culture in MEM medium, it disintegrated tumour spheroids in the same cell culture medium. Secondly, we show that media formulation modulates the sensitivity of MDA-MB-231 cells to metformin in the absence of pyruvate. Based on interaction between pyruvate depletion with depletion of glucose and non-essential amino acids, our results suggest that pyruvate, which can be obtained directly from cell culture media or derived from glucose, serine and glycine, blocks the effects of metformin on MDA-MB-231 cells. Therefore, media formulation as well as cell culture model (monolayer culture vs. tumour spheroids) must be considered, when we evaluate the effects of metformin on MDA-MB-231 cells as a function of nutrient availability.

## Acknowledgements

We thank Dr. Jasna Lojk, Suzana Semič, Gregor Bizjak and Jernej Repas for technical assistance.

## Author Contributions

Conceived and designed the experiments: MB PM MP SP. Performed the experiments: MB PM. Analyzed the data: MB PM MP SP. Wrote the paper: MB MP SP.

